# NeoDTI: Neural integration of neighbor information from a heterogeneous network for discovering new drug-target interactions

**DOI:** 10.1101/261396

**Authors:** Fangping Wan, Lixiang Hong, An Xiao, Tao Jiang, Jianyang Zeng

## Abstract

**Motivation:** Accurately predicting drug-target interactions (DTIs) *in silico* can guide the drug discovery process and thus facilitate drug development. Computational approaches for DTI prediction that adopt the systems biology perspective generally exploit the rationale that the properties of drugs and targets can be characterized by their functional roles in biological networks.

**Results:** Inspired by recent advance of information passing and aggregation techniques that generalize the convolution neural networks (CNNs) to mine large-scale graph data and greatly improve the performance of many network-related prediction tasks, we develop a new nonlinear end-to-end learning model, called NeoDTI, that integrates diverse information from heterogeneous network data and automatically learns topology-preserving representations of drugs and targets to facilitate DTI prediction. The substantial prediction performance improvement over other state-of-the-art DTI prediction methods as well as several novel predicted DTIs with evidence supports from previous studies have demonstrated the superior predictive power of NeoDTI. In addition, NeoDTI is robust against a wide range of choices of hyperparameters and is ready to integrate more drug and target related information (e.g., compound-protein binding affinity data). All these results suggest that NeoDTI can offer a powerful and robust tool for drug development and drug repositioning.

**Availability and implementation:** The source code and data used in NeoDTI are available at: https://github.com/FangpingWan/NeoDTI.

**Supplementary information:** Supplementary data are available at *Bioinformatics* online.

## 1 Introduction

Identifying drug-target interactions (DTIs) through computational approaches can greatly narrow down the large search space of drug candidates for downstream experimental validation, and thus significantly reduce the high cost and the long period of developing a new drug (Langley *et al.*, 2017). Currently, the structure based (Morris *et al.*, 2009), ligand-similarity based (Keiser *et al.*, 2007) and machine learning based methods (Yuan *et al.*, 2016; Luo *et al.*, 2017) are three main classes of prediction approaches in computational aided drug screening. The structure based methods generally require the three-dimensional (3D) structures of proteins and have limited performance for those proteins with unknown structures, which unfortunately is the case for a majority of targets. The ligand-similarity based methods exploit the common knowledge of known interacting ligands to make prediction. Such approaches cannot lead to confident prediction results if the compound of interest is not indicated in the library of reference ligands. Recently, the machine learning based methods (Bleakley and Yamanishi, 2009; Luo *et al.*, 2017), which fully exploit the latent correlations among the related features of drugs and targets have become a highly promising strategy for DTI prediction. For instance, the DTI network data have been integrated with the drug structure and protein sequence information into a network-based machine learning model (e.g., a regularized least squares framework) for predicting new DTIs (Xia *et al.*, 2010; van Laarhoven *et al.*, 2011; van Laarhoven and Marchiori, 2013). Inspired by the recent surge of deep learning techniques, models with higher predictive capacity have also been developed in various drug discovery settings (e.g., compound-protein interaction prediction, drug discovery with one-shot learning) (Wang and Zeng, 2013; Wan and Zeng, 2016; Hamanaka *et al.*, 2017; Tian *et al.*, 2016; Altae-Tran *et al.*, 2017; Xu *et al.*, 2017).

In addition to known DTI data, chemical structure, protein sequence information, and other properties of drugs and targets can also be characterized by their various functional roles in biological systems (e.g., protein-protein interactions and drug-disease associations). Indeed, by integrating diverse information from heterogeneous data sources, methods like DTINet (Luo *et al.*, 2017), MSCMF (Zheng *et al.*, 2013), HNM (Wang *et al.*, 2014) can further improve the accuracy of DTI prediction. However, these methods still suffer from certain limitations that need to be addressed. For example, in MSCMF (Zheng *et al.*, 2013), the employed matrix factorization operation of a given drug-target interaction network is regularized by the corresponding drug and protein similarity matrices, which are obtained by integrating multiple data sources through a weighted averaging scheme. Under such a data integration strategy, substantial loss of information may occur and thus result in a sub-optimal solution. DTINet (Luo *et al.*, 2017) first uses an unsupervised manner to automatically learn low-dimensional feature representations of drugs and targets from heterogeneous network data, and then applies inductive matrix completion (Natarajan and Dhillon, 2014) to predict new DTIs based on the learnt features. In such a framework, separating feature learning from the prediction task at hand may not yield the optimal solution, as the features learnt from the unsupervised learning procedure may not be the most suitable representations of drugs or targets for the final DTI prediction task. In addition, by constraining the learning models to only take relatively simple forms (e.g., bilinear or log-bilinear functions), these methods may not be sufficient enough to capture the complex hidden features behind the heterogeneous data. Recent advance of information passing and aggregation techniques that generalize the conventional convolution neural networks (CNNs) to large-scale graph data have shown substantial performance improvement on the network-related prediction tasks (Hamilton *et al.*, 2017; Gilmer *et al.*, 2017). This inspires us to incorporates deeper learning models to extract complex information from a highly heterogeneous network and discover new DTIs.

In this paper, we propose a new framework, called NeoDTI (**NE**ural integration of neighb**O**r information for **DTI** prediction) to predict new drug-target interactions from heterogeneous data. NeoDTI integrates neighborhood information of the heterogeneous network constructed from diverse data sources via a number of information passing and aggregation operations, which are achieved through the nonlinear feature extraction by neural networks. After that, NeoDTI applies a network topology-preserving learning procedure to enforce the extracted feature representations of drugs and targets to match the observed networks. Comprehensive tests on several challenging and realistic scenarios in DTI prediction have demonstrated that our end-to-end prediction model can significantly outperform several baseline prediction methods. Moreover, several novel DTIs predicted by NeoDTI with evidence supports from previous studies in the literature further indicate the strong predictive power of NeoDTI. In addition, the robustness of NeoDTI and its extendability to integrate more heterogeneous data (e.g., compound-protein binding affinity data) have been examined through various tests. All these results suggest that NeoDTI can provide a powerful and useful tool in predicting unknown DTIs, and thus advance the drug discovery and repositioning fields.

## 2 Methods

### 2.1 Problem formulation

NeoDTI predicts unknown drug-target interactions (DTIs) from a drug and target related heterogeneous network, in which drugs, targets and other objects are represented as nodes, and DTIs and other interactions or associations are represented as edges. We first introduce the definition of a heterogeneous network.

#### DEFINITION 1.

***(Heterogeneous network)***. *A* ***heterogeneous network (HN)*** *is defined as a directed (or undirected) graph G* = (*V, E*), *in which each node v in the node set V belongs to an object type from an object type set O, and each edge e in the edge set E* ⊂ *V* × *V* × *R belongs to a relation type from a relation type set R.*

The datasets used in our framework to construct the HN (also see Section 3.1) include the object type set *O* = {*drug, target, side*-*effect, disease*}, and the relation type set *R* = {*drug*-*structure*-*similarity, drug*-*side*-*effect*-*association, drug*-*protein*-*interaction, drug*- *drug*-*interaction, drug*-*disease*-*association, protein*-*sequence*- *similarity, protein*-*drug*-*interaction, protein*-*disease*-*association, protein*-*protein*-*interaction, disease*-*protein*-*association, disease*-*drug*-*association, side*-*effect*-*drug*-*association*}. In our current framework, each node only belongs to a single object type although it can be relatively easily extended to a multi-object-type mapping scenario. In addition, all edges are undirected and non-negatively weighted. Also, the same two nodes can be linked by more than one edge, e.g., two drugs can be linked by a *drug*-*drug*-*interaction* edge and a *drug*-*structure*-*similarity* edge simultaneously.

Given an HN *G*, NeoDTI aims to automatically learn a network topology-preserving node-level embedding (i.e., a function that maps nodes to their corresponding feature representations that preserve the original topological characteristics as much as possible) from *G* that can be used to greatly facilitate the prediction of drug-target interactions. Most existing techniques for learning the embeddings of structured data mainly exploit the rationale that the elements of these structured data can be well characterized by their contextual information. For example, in natural language processing, the Word2vec technique (Mikolov *et al.*, 2013) enforces the embedding of words to preserve the semantic relationships with their corresponding surrounding words. The graph embedding techniques, such as Deepwalk (Perozzi *et al.*, 2014) and metapath2vec (Dong *et al.*, 2017), have extended this embedding strategy to further learn the latent representations of network data. Recent advance in generalizing convolutional neural networks (CNNs) to analyze large-scale graph data (Defferrard *et al.*, 2016; Kipf and Welling, 2016) and the integration of the information passing and aggregation techniques with different graph convolution operations into a unified framework (Gilmer *et al.*, 2017; Hamilton *et al.*, 2017) have brought significant performance improvement for many network-related prediction tasks, such as predicting the biological activities of small molecules, graph signal processing and social network data analysis. Similar to GraphSAGE (Hamilton *et al.*, 2017) and Message Passing Neural Networks (MPNNs) (Gilmer *et al.*, 2017), our framework NeoDTI also applies neural networks to integrate neighborhood information from individual nodes. However, unlike GraphSAGE which mainly focuses on learning a node-level embedding from a homogeneous network or MPNNs which aim at learning a graph-level embedding from heterogeneous graphs for predicting molecular properties, NeoDTI focuses on learning a node-level embedding from a heterogeneous network. In addition, to the best of our knowledge, NeoDTI is the *first* framework to systematically integrate the neural information passing and aggregation techniques with the topology-preserving optimization scheme into an end-to-end learning framework to extract the latent features of drugs and targets from a heterogeneous network to make DTI prediction.

### 2.2 The worflow of NeoDTI

NeoDTI consists of the following three main steps: (1) neighborhood information aggregation; (2) updating the node embedding; (3) topology-preserving learning of the node embedding. Through Steps (1) and (2), each node in a given HN generates a new feature representation by integrating its neighborhood information with its own features. Through Step (3), we enforce the embedding of nodes to be topology-preserving, which is useful for extracting the topological features of individual nodes for accurate DTI prediction. Next, we will introduce the mathematical formulations of these three steps.

#### Definition 2

(***Neighborhood information aggregation***). *Given an HN G, an initial node embedding function f*^0^ : *V* → ℝ^*d*^ *that maps each node v* ∈ *V to its d-dimensional vector representation f*^0^(*v*) *and an edge weight mapping function s* : *E* → ℝ *that maps each edge e* ∈ *E to a non-negative real value s*(*e*), ***neighborhood information aggregation*** *for node v is defined as:*

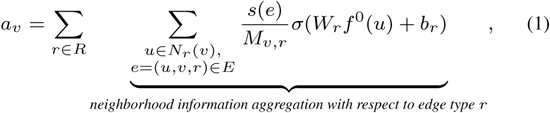

*where N*_*r*_ (*v*) = {*u, u* ∈ *V, u* ≠ *v*, (*u, v, r*) ∈ *E*} *denotes the set of adjacent nodes connected to v* ∈ *V through edges of type r* ∈ *R, σ*(*·*) *stands for a nonlinear activation function over a single-layer neural network parameterized by weights W*_*r*_ ∈ ℝ^*d*×*d*^ *and a bias term b*_*r*_ ∈ ℝ^*d*^, *and 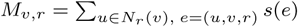 stands for a normalization term.*

More specifically, for each edge type *r*, the neighborhood information aggregation operation for node *v* with respect to *r* can be obtained by first nonlinearly transforming the embedded feature representations of the corresponding adjacent nodes *f*^0^(*u*), *u* ∈ *N*_*r*_ (*v*) through an edge-type specific single-layer neural network that is parameterized by weights *W*_*r*_ ∈ ℝ^*d*×*d*^, a bias term *b*_*r*_ ∈ ℝ^*d*^ and a nonlinear activation function *σ*(*·*), and then averaging by the normalized edge weight, i.e., 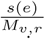. Finally, the output of the neighborhood information aggregation operation *a*_*v*_ for node *v* is the summation of neighborhood information aggregation with respect to every edge type *r*. Here, the initial node embedding *f*^0^(*u*), *∀u* ∈ *V* is obtained through a random mapping.

#### Definition 3

***(Updating the node embedding)***. *Given the aggregated neighbor information a*_*v*_*’s for all nodes v’s, the process of* ***updating the node embedding*** *is defined as:*

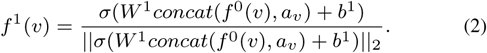

The above equation states that the new embedding of node *f*^1^(*v*) can be obtained using a single-layer neural network that is parameterized by weights *W*^1^ ∈ ℝ^(2*d*)×*d*^, a bias term *b*^1^ ∈ ℝ^*d*^ and a nonlinear activation function *σ*(·) to nonlinearly transform the concatenation of the original embedding *f*^0^(*v*) and the neighborhood aggregation information *a*_*v*_, and then normalized by its *l*_2_ norm.

Noted that in principle we could repeat the previous two steps alternately several times to produce more embeddings of nodes (e.g., *f*^2^(*·*), *f*^3^(·), *…*). In practice, we find that we only need to conduct such a process once to obtain reasonably good prediction results, according to our validation tests (as described in Supplementary Materials). In the rest part of this section, we will mainly use *f*^1^(·) to demonstrate our algorithm for convenience. In addition, we choose to use *ReLU* (*x*) = *max*(0, *x*) as the activation function *σ*(·).

#### Definition 4

***(Topology-preserving learning of the node embedding)***. *Given the embedding of nodes f*^1^(·), ***topology-preserving learning of the node embedding*** *is defined as:*

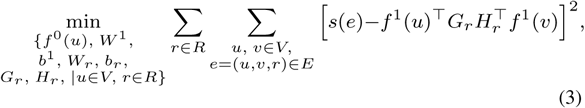

*where G*_*r*_, *H*_*r*_ ∈ ℝ^*d*×*k*^ *are edge-type specific projection matrices.*

The above equation states that, after edge-specific projections of *f*^1^(*u*) and *f*^1^(*v*) by *G*_*r*_ and *H*_*r*_, respectively, the inner product of the two projected vectors should reconstruct the original edge weight *s*(*e*) as much as possible. Note that a similar reconstruction strategy has also been used in (Luo *et al.*, 2017; Natarajan and Dhillon, 2014) to solve the link prediction problems. In addition, if the edge type *r* is symmetric, i.e., *r* ∈ {*drug*-*structure*-*similarity, protein*-*sequence*-*similarity, drug*-*drug*-*interaction, protein*-*protein*-*interaction*}, we use the tie weights (i.e., *G*_*r*_ = *H*_*r*_) to enforce this symmetric property. Here, the summation of the squared reconstruction errors is minimized for all edges with respect to all unknown parameters. Since all mathematical operations in Equations 1, 2 and 3 are differentiable or subdifferentiable (e.g., for the ReLU activation function), all parameters can be trained through an end-to-end manner by performing gradient descent to minimize the final objective function described in Equation 3.

Finally, after Step (3), the predicted interaction confidence score between drug node *u* and protein node *v* can be obtained by

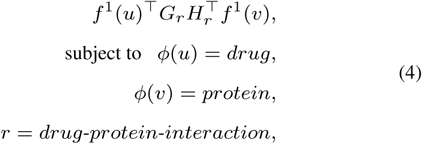

where *ϕ*(*u*) and *ϕ*(*u*) stand for the node types of *u* and *v*, respectively, and *r* represents their edge type.

The above operation is equivalent to reconstructing the *drug*-*protein* edge weight between nodes *u* and *v*. By collecting *f*^1^(*u*)’s for all drugs and *f*^1^(*v*)’s for all targets, we can form a drug feature matrix *F*_*drug*_ and a target feature matrix *F*_*target*_. Then, the reconstructed drug-target interaction matrix can be written as:

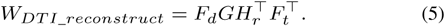

In this sense, we can consider our DTI prediction task as a matrix factorization or completion problem. However, unlike the conventional matrix factorization approaches (Zheng *et al.*, 2013; Natarajan and Dhillon, 2014), NeoDTI incorporates a deeper learning model to construct the feature matrices *F*_*d*_ and *F*_*t*_ by explicitly defining the construction processes of *F*_*d*_ and *F*_*t*_ through Steps (1) and (2). In addition, through these two steps, NeoDTI incorporates the prior knowledge of network topology into *F*_*d*_ and *F*_*t*_ and specifies the forms of these two matrices to guide the downstream optimization process. As a result, NeoDTI prevents the DTI network as well as other networks from being factorized arbitrarily in Step (3), which can serve as a useful regularizer and thus lead to performance improvement for DTI prediction (as also demonstrated in our cross-validation tests; see the Results section).

## 3 Results

### 3.1 Datasets

We adopted the datasets that were curated in our previous study (Luo *et al.*, 2017), which included six individual drug/protein related networks: **drug-protein interaction** and **drug-drug interaction** networks (interactions were extracted from Drugbank Version 3.0 (Knox *et al.*, 2010)), the **protein-protein interaction** network (interactions were extracted from the HPRD database Release 9 (Keshava Prasad *et al.*, 2008)), **drug-disease association** and **protein-disease association** networks (associations were extracted from the Comparative Toxicogenomics Database (Davis *et al.*, 2012)) and the **drug-side-effect association** network (associations were extracted from the SIDER database Version 2 (Kuhn *et al.*, 2010)). The basic statistics of these datasets can be found in Table S1 in Supplementary Materials. We also incorporated drug chemical structure information as well as protein sequence information by creating two extra networks: the **drug structure similarity** network (i.e., a pair-wise chemical structure similarity network measured by the dice similarities of the Morgan fingerprints with radius 2 (Rogers and Hahn, 2010), which were computed by RDKit (Landrum, 2006)) and the **protein sequence similarity** network (which was obtained based on the pair-wise Smith-Waterman scores (Smith and Waterman, 1981)). All networks had binary edge weights (one represents a known interaction or association, and zero otherwise) except the drug structure similarity and the protein sequence similarity networks, which had non-negative real-valued edge weights. We combined all these eight networks to construct the heterogeneous network (Figure 1) for evaluating the prediction performance of NeoDTI.

**Fig. 1.**
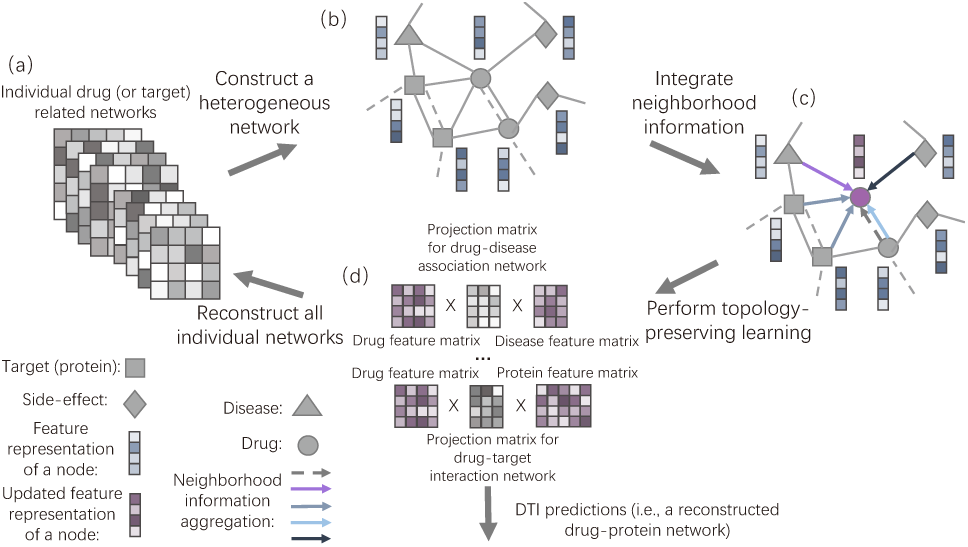
The schematic workflow of NeoDTI. (a) NeoDTI uses eight individual drug or target related networks (see Section 3.1 for more details of the used datasets). (b) NeoDTI first constructs a heterogeneous network from these eight networks. Different types of nodes are connected by distinct types of edges. Two nodes can be connected by more than one edge (e.g., a solid link representing drug-drug-interaction and a dashed link representing drug-structure-similarity). In addition, NeoDTI associates each node with a feature representation. (c) To extract information from neighborhood, each node adopts a neighborhood information aggregation operation (see Definition 2 in the main text). Each colored arrow represents a specific aggregation function with respect to a specific edge type. Then each node updates its feature representation by integrating its current representation with the aggregated information (see Definition 3 in the main text). (d) By enforcing the node features to reconstruct the original individual networks as much as possible (see Definition 4 in the main text), NeoDTI effectively learns the topology-preserving node features that are useful for drug-target interaction prediction.

### 3.2 NeoDTI yields superior performance in predicting new drug-target interactions

The DTI prediction can be considered as a binary classification problem, in which the known interacting drug-target pairs are regarded as positive examples, while the unknown interacting pairs are treated as negative examples. Several challenging and realistic scenarios were considered in our tests to evaluate the prediction performance of NeoDTI. The hyperparameters of NeoDTI were determined using an independent validation set (as described in Supplementary Materials). We first ran a ten-fold cross validation test on all positive pairs and a set of randomly sampled negative pairs, whose number was ten times as many as that of positive samples. This scenario basically mimicked the practical situation in which the drug-target interactions are sparsely labeled. For each fold, a randomly chosen subset of 90% positive and negative pairs was used as training data to construct the heterogeneous network and then train the parameters of NeoDTI (i.e., during the topology-preserving learning process, we only calculated the reconstruction loss of the DTI network with respect to training data, while the reconstruction losses of other types of networks were computed as usual), and the remaining 10% positive and negative pairs were held out as the test set. We also compared the performance of NeoDTI with that of five baseline methods, including DTINet (Luo *et al.*, 2017), HNM (Wang *et al.*, 2014), NetLapRLS (Xia *et al.*, 2010), DT-Hybrid (Alaimo *et al.*, 2013) and BLMNII (Mei *et al.*, 2012). The area under precision recall (AUPR) curve and the area under receiver operating characteristic (AUROC) curve were used to evaluate prediction performance of all prediction methods. We observed that NeoDTI greatly outperformed other baseline methods, with significant improvement (6.5% in terms of AUPR and 2.7% in terms of AUROC) over the second best method (Figures 2a and S1a).

**Fig. 2.**
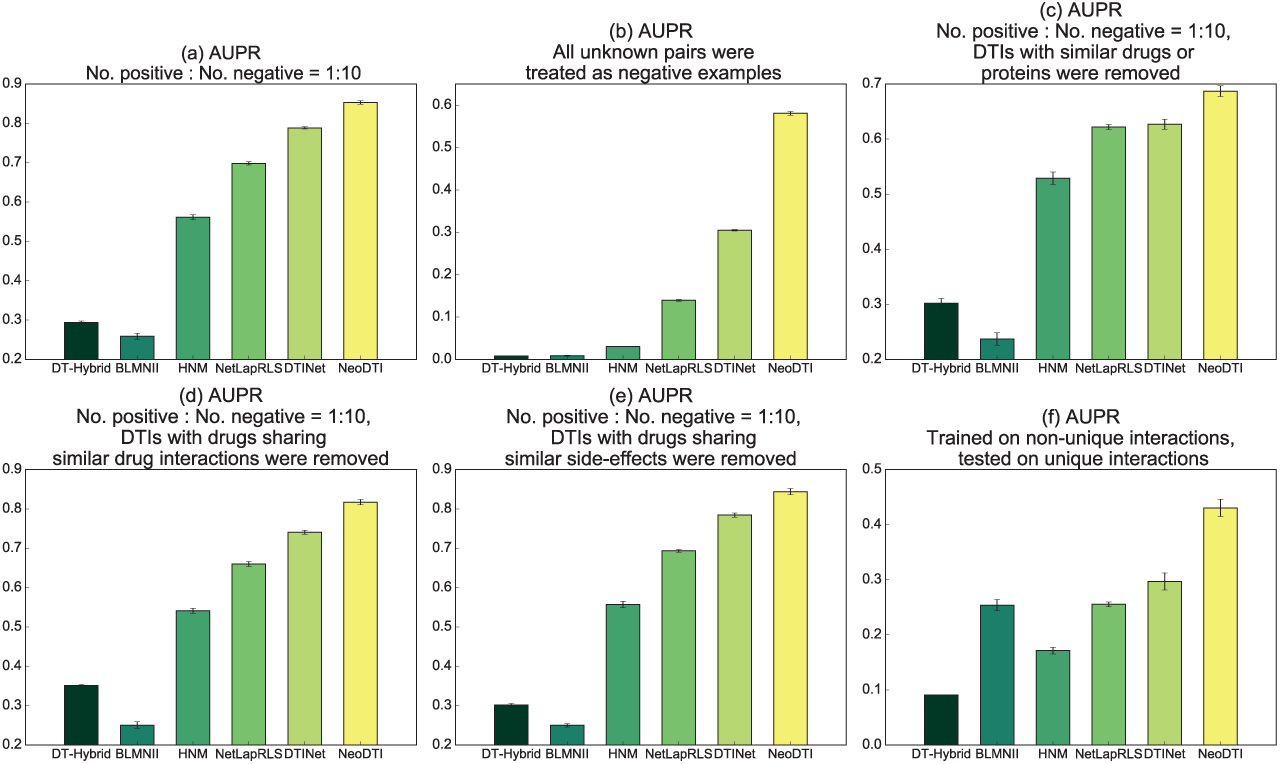
Performance evaluation of NeoDTI on several challenging scenarios in terms of the AUPR scores. (a) A ten-fold cross-validation test in which the ratio between positive and negative samples was set to 1 : 10. (b) A ten-fold cross-validation test in which all unknown drug-target interacting pairs were considered. (c-e) Ten-fold cross-validation with positive : negative ratios = 1 : 10 on several scenarios of removing redundancy in data: (c) DTIs with similar drugs and proteins were removed; (d) DTIs with drugs sharing similar drug interactions were removed; (e) DTIs with drugs sharing similar side-effects were removed. (f) NeoDTI was trained on non-unique drug-target interacting pairs and tested on unique drug-target interacting pairs. More details on the baseline methods can be found in Supplementary Materials. All results were summarized over 10 trials and expressed as mean *±* standard deviation.

Next, we further increased the positive-negative ratio by including all negative examples (i.e., all unknown drug-target interacting pairs) in the ten-fold cross-validation procedure (the ratio between positive and negative samples was around 1.8 × 10^−3^). We observed a larger AUPR improvement (27.6%) over the second best method (Figure 2b). Although NeoDTI, DTINet and NetLapRLS achieved comparable results in terms of AUROC in this scenario (Figure S1b), as also stated in previous work (Davis and Goadrich, 2006), here, AUPR generally provides a more informative criterion than AUROC for the highly skewed datasets. Since drug discovery is generally a needle-in-a-haystack problem, the substantial improvement in AUPR truely demonstrated the superior prediction performance of NeoDTI over other methods.

Since the datasets may contain “redundant” DTIs (i.e., a same protein is connected to more than one similar drugs and vice versa), the prediction performance can be easily inflated by easy predictions in this case (Luo *et al.*, 2017). To consider this issue, we followed the same evaluation strategies as in (Luo *et al.*, 2017) by conducting the following additional ten-fold cross validation tests: (1) removing DTIs with similar drugs (i.e., drug chemical structure similarities *>* 0.6) or similar proteins (i.e., protein sequence similarities *>* 40%); (2) removing DTIs with drugs sharing similar drug interactions (i.e., Jaccard similarities *>* 0.6); (3) removing DTIs with drugs sharing similar side-effects (i.e., Jaccard similarities *>* 0.6); (4) removing DTIs with drugs or proteins sharing similar diseases (i.e., Jaccard similarities *>* 0.6). In all these test scenarios, we kept the ratios between positive and negative samples to be 1 : 10. As expected, we observed a drop in prediction performance for all prediction methods after the removal of redundant DTIs (Figures 2c-e, S1c-g). However, NeoDTI still consistently outperformed other prediction methods in terms of both AUPR and AUORC, which also indicated the robustness of NeoDTI after removing the redundancy in data.

In dyadic prediction, if a dataset contains many drugs or targets with only one interacting partner, conventional cross-validation may not be a proper way to evaluate the prediction performance. Here, we call such drugs, proteins and interactions as “unique”. In such a case, conventional training methods may lean to exploit the bias towards those unique drugs and targets to boost the performance (van Laarhoven and Marchiori, 2014). To investigate this issue, we further evaluated the prediction performance of NeoDTI by separating unique DTIs from non-unique ones. That is, all methods were trained on non-unique DTIs and then evaluated on unique DTIs. Note that in such a case, the negative examples in the test data were sampled by enforcing the corresponding drugs or targets (or both) to be unique. This scenario basically mimicked the situation in which the DTIs of new drugs or targets are predicted without much prior DTI knowledge.

We found that NeoDTI significantly outperformed all the baseline methods at least by 13.3% in terms of AUPR, which suggested that NeoDTI can have a much better generalization capacity over other state-of-the-art methods, when predicting new DTIs for those drugs or targets without much prior DTI knowledge.

### 3.3 Robustness of NeoDTI

In this section, we further evaluated the robustness of NeoDTI by varying different types of data used in the heterogeneous network as well as the hyperparameters of NeoDTI. All computational experiments in this section were conducted using a ten-fold cross-validation procedure in which the ratios of positive versus negative samples were set to 1 : 10.

To examine the effects of incorporating heterogeneous data, we first evaluated the performance of NeoDTI when being trained using only the drug-protein interaction network. We observed a substantial drop of prediction performance (11.1% in terms of AUPR and 9% in terms of AUROC), compared to that of the original NeoDTI model trained on all eight networks (Figure 3a). We then investigated the effects of incorporating individual networks by training NeoDTI again on a heterogeneous network constructed from each individual network and the drug-protein interaction network. As expected, we found that adding individual drug or target related networks can improve the prediction performance (Figures 3a, S2a-e). These results suggested that diverse information from multiple data sources can better characterize the latent properties of drugs and targets, and thus incorporating heterogeneous information is necessary to improve the accuracy of DTI prediction. In addition, to examine whether NeoDTI can also be easily extended to incorporate more drug or target related information beyond the previously used datasets, we further incorporated compound-protein binding affinity information into the heterogeneous network. More specifically, we collected all the binding affinity data between drug-like compounds and proteins that satisfied *Ki* ≤ 1.0*nm* from the ZINC15 database (Sterling and Irwin, 2015). In total, we extracted 1,696 edges that connected 1, 244 compounds to the proteins used in our previous datasets. We set the negative logarithm of *Ki* between a pair of compound and protein as the weight of their corresponding interaction. We also linked the compounds and drugs by drug-structure-similarity edges. In addition, if a compound and a drug were similar (i.e., chemical structure similarities *>* 0.6) and connected to the same protein in the test set, we removed this compound-protein pair from the training set to ease the inflation of prediction performance that may be resulted from the redundancy in data. We found that NeoDTI trained on this new heterogeneous network further improved AUPR from 85.3% to 86.2% and AUROC from 94.6% to 95.1% (Figure 3b), which demonstrated the easy extendability of NeoDTI to integrate more heterogeneous information.

**Fig. 3.**
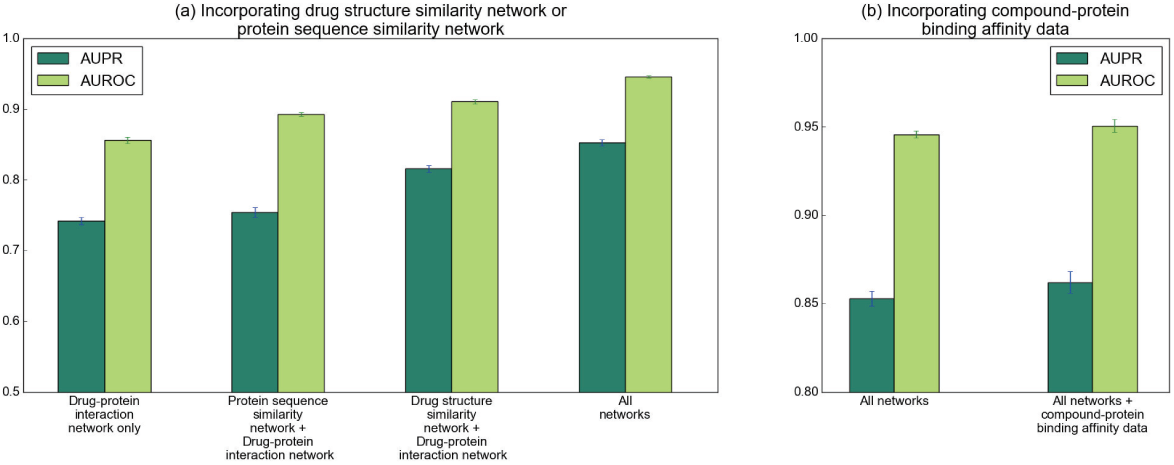
Incorporating more drug or target related information can improve the prediction performance of NeoDTI. (a) Incorporating the drug structure similarity network or protein sequence similarity network. (b) Incorporating the compound-protein binding affinity data. All results were summarized over 10 trials and expressed as mean *±* standard deviation.

In our topology-preserving learning of the node embedding, we enforce the feature representations of nodes to reconstruct all types of edges as much as possible. We further investigated the effect of this edge reconstruction strategy by conducting an additional test in which we constrained NeoDTI to only reconstruct the DTI edges. In this test, we observed a decrease in prediction performance with 5.5% in terms of AUPR and 2.7% in terms of AUROC (Figure S2f). Thus, reconstructing other types of network edges is useful for boosting the prediction performance. Such an operation probably serves as a beneficial regularizer to further overcome the potential overfitting problem.

In addition, we investigated the robustness of NeoDTI against different choices of hyperparameters: (1) For the dimension *d* of the node embedding, we tested *d* = 256, 512 and 1024; (2) For the dimension *k* of the projection matrices, we tested *k* = 256, 512 and 1024; (3) For the repetition time *p* of neighborhood information aggregation, we examined *p* = 0, 1, 2 and 3. We found that NeoDTI can produce relatively stable results over a wide range of choices for both *d* and *k*, although we observed that increasing the value of *d* can slightly improve the prediction results (Figure S3a-b). More importantly, we observed significant performance improvement when *p ≥* 1, demonstrating the necessity of integrating neighborhood information for the representation learning of node features (Figure S3c). However, we found that increasing the repetition time of neighborhood information aggregation from one to three did not improve the prediction performance (Figure S3c). Thus, in practice, we only need to run the operation of integrating neighborhood information once.

### 3.4 NeoDTI reveals novel DTIs with literature supports

We also predicted the novel DTIs by training NeoDTI using the whole heterogeneous network, including the aforementioned binding affinity data. We excluded those easy predictions by removing the predicted DTIs that were similar to the known DTIs (i.e., drug chemical structure similarities *>* 0.6 and protein sequence similarities *>* 40%). We then analyzed the predicted DTIs whose prediction confidence scores were significant (three-sigma rule) with respect to the corresponding drugs and targets. The network visualization of the top 100 novel drug-target interactions predicted by NeoDTI can be found in Figure 4.

**Fig. 4.**
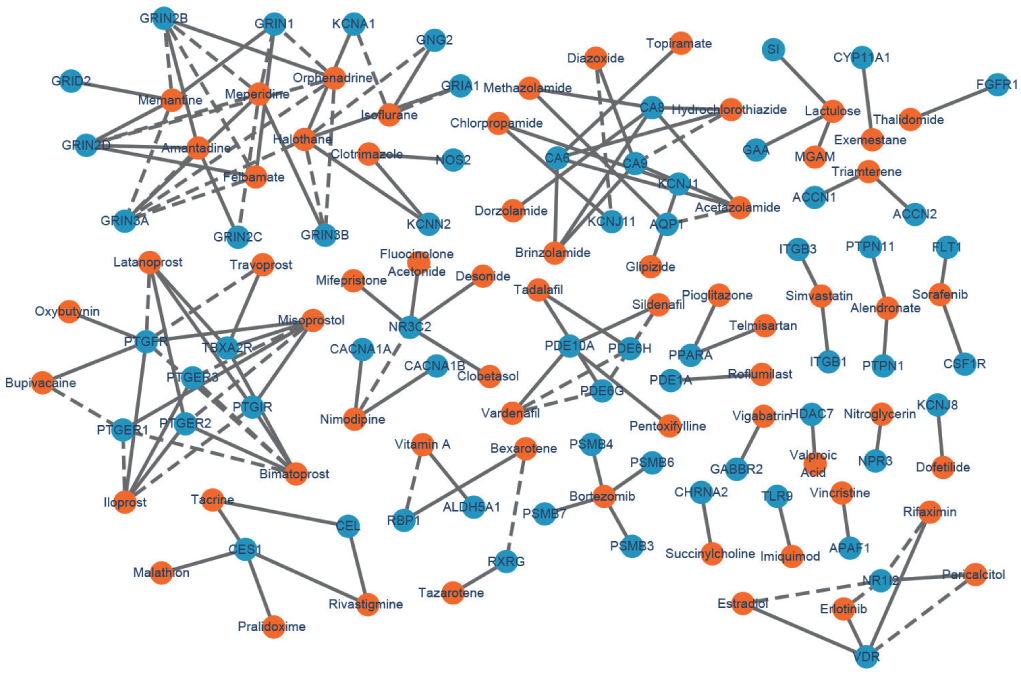
Network visualization of the top 100 novel drug-target interactions predicted by NeoDTI. Blue and orange nodes represent proteins and drugs, respectively. Dashed and solid lines represent the known and predicted drug-target interactions, respectively.

Among the top twenty predicted DTIs ranked according to their confidence scores, eight DTIs can be supported by previous studies in the literature (Table S2). For instance, sorafenib, a drug previously approved for the treatment of advanced renal cell carcinoma, was predicted by NeoDTI to interact with the colony stimulating factor 1 receptor (CSF1R), which plays an important role in the development of mammary gland and mammary gland carcinogenesis (Tamimi *et al.*, 2008). Such a prediction can be supported by a previous study indicating that sorafenib can block CSF1R and induce apoptosis in various classical Hodgkin lymphoma cell lines (Ullrich *et al.*, 2011). In addition, the carbonic anhydrase 6 (CA6), an enzyme abundantly found in salivary glands, has been previously reported to be the target of three drugs, including zonisamide, ellagic acid and mafenide (Knox *et al.*, 2010), was predicted by NeoDTI to also interact with acetazolamide. This prediction can be supported by the previous finding on the CA6 inhibitory activity of acetazolamide (Nishimori *et al.*, 2007). Overall, these novel DTIs predicted by NeoDTI with literature supports further demonstrated its strong predictive power.

## 4 Conclusion

In this paper, we develop a new framework, called NeoDTI, to integrate diverse information from a heterogeneous network to predict new drug-target interactions. NeoDTI extracts the complex hidden features of drugs and targets by applying neural networks to integrate neighborhood information in the input heterogeneous network. By simultaneously optimizing the feature extraction process and the DTI prediction model through an end-to-end manner, NeoDTI can achieve superior prediction performance over other state-of-the-art methods. The effectiveness and robustness of NeoDTI have been extensively validated on several realistic prediction scenarios and supported by the finding that many of the novel predicted DTIs agree well with the previous studies in the literature. Moreover, NeoDTI can incorporate more drug and target related information readily (e.g., compound-protein binding affinity data). Therefore, we believe that NeoDTI can provide a powerful and useful tool to facilitate the drug discovery and drug repositioning processes. In the future, we will further extend NeoDTI by integrating more heterogeneous information and validate some of the prediction results through wet-lab experiments.

## Acknowledgements

The authors thank Yunan Luo and Hailin Hu for helpful discussions.

