## Supplementary Materials for "NeoDTI: Neural integration of neighbor information from a heterogeneous network for discovering new drug-target interactions"

### 1 Hyperparameter Selection

We used an independent validation dataset to determine the values of hyperparameters, including the dimension $d \in \{256, 512, 1024\}$ of node embedding, the dimension $k \in \{256, 512, 1024\}$ of the edge-type specific projection matrices, the repetition time $p \in \{0, 1, 2, 3\}$ for alternately repeating Steps (1) and (2), the gradient clipping norm from $\{1, 5\}$ and the number of steps for performing gradient descent. In particular, after splitting the data into training and test datasets in the cross-validation procedure, we randomly separated 0.5% of the training set as an independent validation dataset. We tuned the above hyperparameters based on this separate validation set and evaluated the performance of NeoDTI on the test dataset. In addition, we used the Adam optimizer [1] with the default learning rate 0.001 to perform gradient descent.

### 2 Baseline Methods

We compared the performance of NeoDTI with that of several network-based DTI prediction methods, including DT-Hybrid [2], BLMNII [3], HNM [4], NetLapRLS [5] and DTINet [6]. DT-Hybrid [2], BLMNII [3] and LPMIHN [7] have been reported as the state-of-the-art network-based prediction methods [8]. LPMIHN was not included in our comparisons because we encountered technical difficulty in implementing this method. Among the baseline methods used in our comparison tests, DTINet and HNM can integrate multiple heterogeneous information to predict new DTIs, while the other methods are not particularly designed to exploit multiple drug or protein network data for DTI prediction. To make a fair comparison, we followed the same strategy as in [6] to integrate multiple networks into a single network for DT-Hybrid, BLMNII and NetLapRLS in our comparison tests. In particular, for all interaction or association networks, i.e., drug-drug interaction, drug-disease association, drug-side-effect association, protein-protein interaction and protein-disease association networks, we constructed the corresponding drug-drug or protein-protein similarity networks based on the Jaccard similarities. Then, the final similarity score between drugs $i$ and $j$ after integrating all similarity networks was obtained by $1 - \prod_k (1 - d_{ij}^k)$, where $d_{ij}^k \in [0, 1]$ stands for the similarity between drugs $i$ and $j$ based on the network $k$. Here, $k$ can stand for a drug-drug interaction, drug-disease association, drug-side-effect association or drug-structure-similarity network. Similarly, the final similarity score between proteins $i$ and $j$ after integrating all similarity networks was obtained by $1 - \prod_k (1 - p_{ij}^k)$, where $p_{ij}^k \in [0, 1]$ stands for the similarity between proteins $i$ and $j$ based on the network $k$. Here, $k$ can stand for a protein-protein interaction, protein-disease association or protein-sequence-similarity network. Note that the edge weights in the protein-sequence-similarity network are ranged from $[0, 100]$. Here, we normalized the weights to $[0, 1]$. We used the above final similarity score as edge weights to construct the drug-drug and protein-protein similarity networks, and used them in the baseline methods DT-Hybrid, BLMNII and NetLapRLS. For the hyperparameters in all baseline methods, we used the default settings as reported in the original papers.

### 3 Supplementary Figures

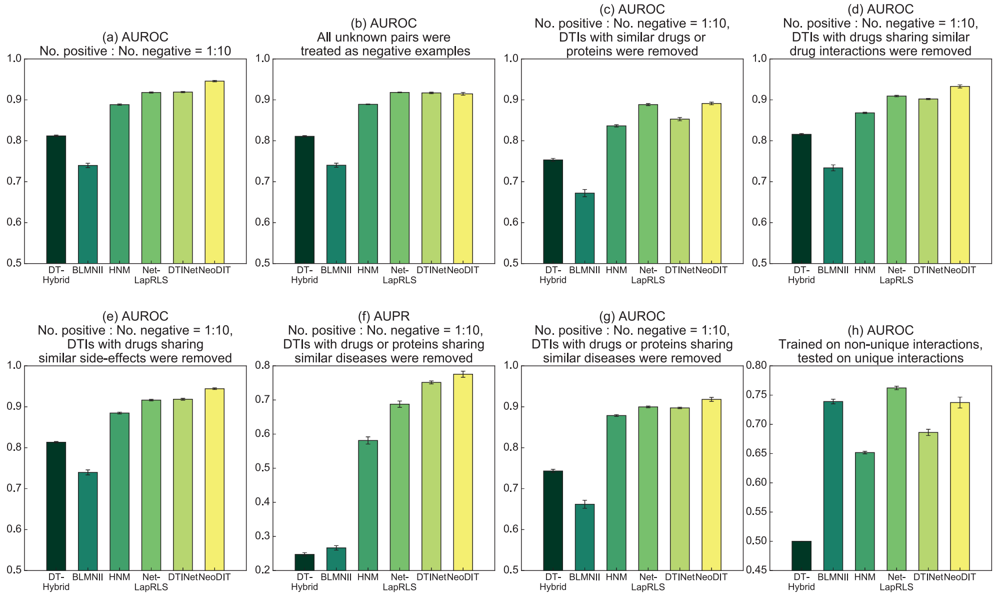


Figure S1. Supplementary results on the performance evaluation of NeoDTI on several challenging scenarios in terms of the AUPR and AUROC scores. (a) A ten-fold cross-validation test in which the ratio between positive and negative samples was set to 1 : 10. (b) A ten-fold cross-validation test in which all unknown drug-target interacting pairs were considered. (c-g) Ten-fold cross-validation with positive : negative ratios = 1 : 10 on several scenarios of removing redundancy in data: (c) DTIs with similar drugs and proteins were removed. (d) DTIs with drugs sharing similar drug interactions were removed. (e) DTIs with drugs sharing similar side-effects were removed. (f,g) DTIs with drugs and proteins sharing similar diseases were removed. (h) NeoDTI was trained on non-unique drug-target interacting pairs and tested on unique drug-target interacting pairs. More details on the baseline methods can be found in this Supplementary Materials. All results were summarized over 10 trials and expressed as mean ± standard deviation.

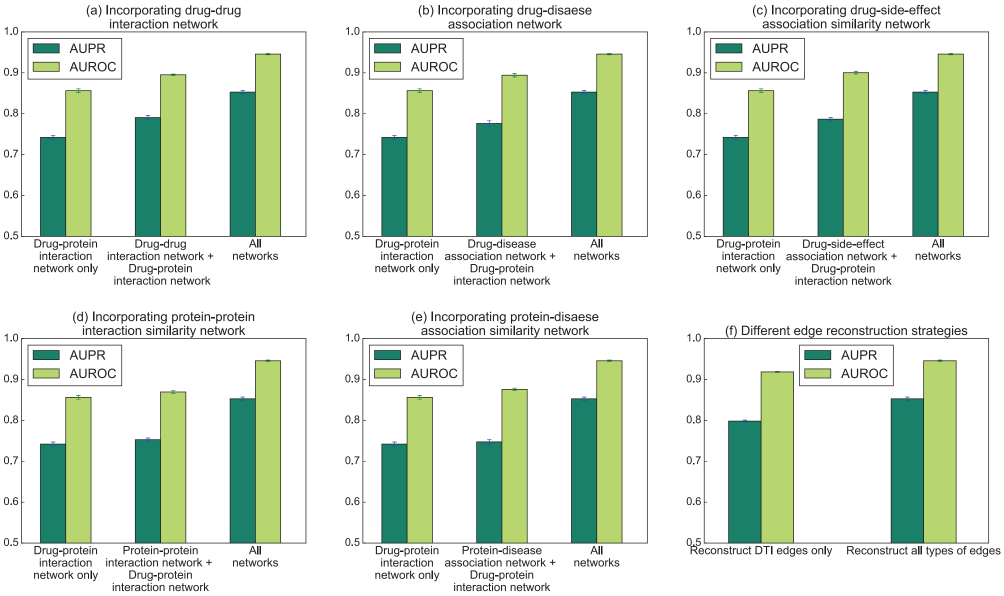


Figure S2. Supplementary results on the effects of incorporating heterogeneous information in NeoDTI. (a) Incorporating the drug-drug interaction network. (b) Incorporating the drug-disease association network. (c) Incorporating the drug-side-effect association network. (d) Incorporating the protein-protein interaction network. (e) Incorporating the protein-disease association network. (f) Performance under different edge reconstruction strategies. All results were summarized over 10 trials and expressed as mean ± standard deviation.

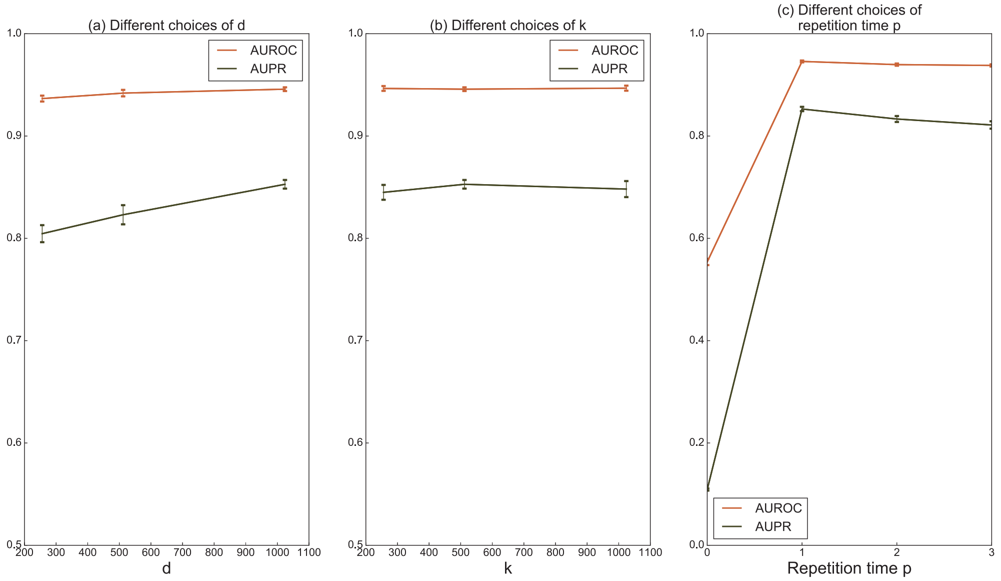


Figure S3. The robustness of NeoDTI over different choices of hyperparameters. (a) Performance of NeoDTI under different choices of the embedding dimension $d$. (b) Performance of NeoDTI under different choices of the dimension $k$ of the projection matrices. (c) Performance of NeoDTI under different choices of repetition time $p$ for performing neighborhood information integration. All results were summarized over 10 trials and expressed as mean $\pm$ standard deviation.

### 4 Supplementary Tables

| Node | Count |
|---|---|
| Drug | 708 |
| Protein | 1,512 |
| Disease | 5,603 |
| Side-effect | 4,192 |
| Total | 12,015 |

(a)

| Edge | Count |
|---|---|
| Drug-Protein | 1,923 |
| Drug-Drug | 10,036 |
| Drug-Disease | 199,214 |
| Drug-Side-effect | 80,164 |
| Protein-Protein | 7,363 |
| Protein-Disease | 1,596,745 |
| Total | 1,895,445 |

(b)

Table S1. (a) The node statistics and (b) the edge statistics of the datasets used in our computational experiments. The datasets were curated in our previous study [6].

| Drug name | Protein name | Supporting references |
|---|---|---|
| Sorafenib | FLT1 | [9] |
| Mifepristone | NR3C2 | [10, 11] |
| Tazarotene | RXRG | [12] |
| Felbamate | GRIN2D | [13] |
| Acetazolamide | CA6 | [14, 15] |
| Rivastigmine | CES1 | [16] |
| Pioglitazone | PPARA | [17] |
| Sorafenib | CSF1R | [18] |

Table S2. Eight novel drug-target interactions among the list of top 20 significant predictions derived by NeoDTI that can be supported by previous studies in the literature.

[5] Zheng Xia, Ling-Yun Wu, Xiaobo Zhou, and Stephen TC Wong. Semi-supervised drug-protein interaction prediction from heterogeneous biological spaces. In *BMC systems biology*, volume 4, page S6. BioMed Central, 2010.

[6] Yunan Luo, Xinbin Zhao, Jingtian Zhou, Jinglin Yang, Yanqing Zhang, Wenhua Kuang, Jian Peng, Ligong Chen, and Jianyang Zeng. A network integration approach for drug-target interaction prediction and computational drug repositioning from heterogeneous information. *Nature Communications*, 8(1), 2017.

[7] Xiao-Ying Yan, Shao-Wu Zhang, and Song-Yao Zhang. Prediction of drug–target interaction by label propagation with mutual interaction information derived from heterogeneous network. *Molecular BioSystems*, 12(2):520–531, 2016.

[8] Maryam Lotfi Shahreza, Nasser Ghadiri, Sayed Rasoul Mousavi, Jaleh Varshosaz, and James R Green. A review of network-based approaches to drug repositioning. *Briefings in bioinformatics*, page bbx017, 2017.

[9] D Kitagawa, K Yokota, M Gouda, Y Narumi, H Ohmoto, E Nishiwaki, K Akita, and Y Kirii. Activity-based kinase profiling of approved tyrosine kinase inhibitors. *Genes to Cells*, 18(2):110–122, 2013.

[10] David Wishart, David Arndt, Allison Pon, Tanvir Sajed, An Chi Guo, Yannick Djoumbou, Craig Knox, Michael Wilson, Yongjie Liang, and Jason Grant. T3db: the toxic exposome database. *Nucleic Acids Research*, 43(Database issue):928–34, 2015.

[11] Lim Emilia, Pon Allison, Djoumbou Yannick, Knox Craig, Shrivastava Savita, Guo An Chi, Neveu Vanessa, and David S Wishart. T3db: a comprehensively annotated database of common toxins and their targets. *Nucleic Acids Research*, 38(Database issue):D781, 2010.
